# SOX2-VSX2 Co-Occupancy Shapes Retinal Neurogenesis Through Dynamic Chromatin Regulation

**DOI:** 10.1101/2025.05.19.654956

**Authors:** Fuyun Bian, Kimiasadat Golestaneh, Emily Davis, Abdullah Khan, Marwa Daghsni, Keevon Flohr, Silvia Liu, Susana da Silva, Len Pennacchio, Issam Aldiri

## Abstract

Retinal neurogenesis is mediated by the coordinated activities of a complex gene regulatory network (GRN) of transcription factors (TFs) in multipotent retinal progenitor cells (RPCs). How this GRN mechanistically guides neural competence remains poorly understood. In this study, we present integrated transcriptional, genetic, and genomic analyses to uncover the regulatory mechanisms of SOX2, a key factor in establishing neural identity in RPCs. We show that SOX2 is preferentially enriched in the RPC-specific enhancer landscape associated with essential regulators of retinogenesis. Disruption of SOX2 expression impairs retinogenesis, marked by a selective loss of enhancer activity near genes essential for RPC proliferation and lineage specification. We identified the RPC transcription factor VSX2 as a binding partner for SOX2, and together, SOX2 and VSX2 co-target a core, retina-specific chromatin repertoire characterized by enhanced TF binding and robust chromatin accessibility. This cooperative binding establishes a shared SOX2-VSX2 transcriptional code that promotes the expression of critical regulators of neurogenesis while repressing the acquisition of alternative lineage cell fate. Our data illuminate fundamental biological insights on how transcription factors act in concert to drive chromatin-based genetic programs underlying retinal neural identity.

## Introduction

Retinogenesis initiates when multipotent retinal progenitor cells (RPCs) cease proliferation and produce six neuronal cell types and one type of glial cell [1, 2]. The gene regulatory network underlying retinal neurogenesis has been extensively studied, revealing numerous transcription factors that orchestrate the step-wise spatial and temporal execution of genetic programs that lead to the specification and terminal differentiation of competent RPCs [3, 4]. Evidence indicates that the RPC genetic architecture is dominated by a small group of the many hundreds of TFs, establishing an interacting regulatory network that drives the RPC transcriptional programs via feedback and feedforward loops [5–7]. This core group of RPC genes includes pivotal factors such as *Pax6*, *Six3*, *Vsx2*, *Lhx2*, *Sox2*, and *Rax* [8–13]. Genetic studies indicate that mutations in these TFs are associated with developmental ocular diseases such as anophthalmia and microphthalmia (AM)[14–16]. Functional studies in animal models further elucidated the roles of these factors, demonstrating that disruption of the RPC GRN members results in defects in key retinal developmental transitions [12, 17–19].

Evidence suggests that the ability of multipotent progenitor cells to generate a particular cell fate in response to a spatial- or temporal-identity factor is encoded by the enhancer landscape, which is modulated by the recruitment of TFs [20–23]. Yet, many of the RPC TFs are also expressed in various parts of the developing nervous system, raising interesting questions on how such TFs promote highly tissue- and stage-specific transcriptional output. A first step in enhancer activation is to gain accessibility around regulatory elements controlling key genes, which is mediated by a special class of pioneer TFs that has the ability to bind and open closed chromatin [24–26].

Sox2, a SOXB1-HMG box transcription factor, is a key member of the regulatory module that coordinates retinogenesis [27, 28]. Sox2 is expressed in RPCs and is required for their neural competency as mouse conditional knockouts result in decreased proliferation and a delay in the onset of proneural TF expression [11, 29]. Sox2 is also expressed in postmitotic retina in a subset of amacrine cells and in Müller glia where it regulates Müller glia quiescence [30]. Importantly, genetic studies revealed that 10-15% of patients with AM have mutations in *SOX2*, underscoring its critical role in human eye development [31, 32]. SOX2 recognizes specific DNA consensus sequences through its highly conserved HMG domain, and can act as a transcriptional repressor and activator, but limited experimental evidence exists to test these functional properties in the developing retina [10, 27].

Here we show that SOX2 controls neural competence by shaping the active enhancer landscape in RPCs. Conditional Knockout of *Sox2* in RPCs leads to loss of chromatin accessibility underlying enhancer activities. SOX2 binds VSX2 and shares common epigenetic and transcriptional programs that drive retinal proliferation and differentiation. Our data highlight how SOX2 governs chromatin accessibility changes essential for cell fate decisions and underscore a SOX2-VSX2 interaction that establishes a cell-type-specific regulatory environment within RPCs, reinforcing the enhancer landscape specific to retinal differentiation.

## Results

### Sox2 sustains active RPC enhancer landscape associated with neural competence

To investigate the functional impact of loss of *Sox2* on chromatin accessibility, we generated an RPC-specific conditional knockout (*Sox2* cKO) by crossing *Sox2*-floxed mice (JAX Lab) with *Chx10-Cre* mice, which induce Cre-mediated recombination in RPCs [33]. *Sox2* homozygous null mice are viable and fertile but display microphthalmia and optic nerve atrophy, in agreement with prior reports [11, 34] (S1 Fig). Histological analysis of the knockout retinae at E14.5 retinae using the RGC differentiation markers OC2 and TUBB3 confirmed loss of neural potentials (S1 Fig). We profiled the chromatin accessibility of *Sox2* cKO and control retinae at E14.5 using Assay for Transposase-Accessible Chromatin followed by Sequencing (ATAC-Seq) (Fig. 1A-D). We identified 2376 differentially accessible regions (DARs) of which 1794 (75%) are reduced and 582 (25%) are increased (FDR < 0.05, DESeq2; Fig. 1A; Dataset S1). The majority (>90%) of DARs located outside promoter regions, suggesting a selective targeting of enhancers (Fig. 1D). To assess this on a deeper level, we annotated the chromatin signature of DARs using ChromHMM enhancer states of the developing retina [35]. An enhancer state was defined by the co-occurrence of the chromatin marks H3K27Ac, BRD4 and H3K4me1, and a low level of the active promoter mark H3K4me3[35]. We found that all the reduced DARs and 75.9% (442/582) of the increased DARs were marked by enhancer activities. Gene Ontology (GO) analysis using Genomic Regions Enrichment of Annotations Tool (GREAT) [36] indicated that reduced chromatin accessibility was highly associated with functions related to eye development and neuron differentiation whereas increased DARs were related to genes involved in signaling pathways (Fig. 1C). Importantly, motif scanning using hypergeometric optimization of motif enrichment (HOMER)[37] denotes that the reduced DARs were highly enriched with SOX recognition sequences, suggesting a direct involvement of SOX2 in regulating chromatin accessibility of those sites (Fig. 1E). On the contrary, increased DARs were primarily enriched with homeodomain recognition motifs and thus are likely to be indirectly regulated by SOX2 (Fig. 1E).

**Figure 1.**
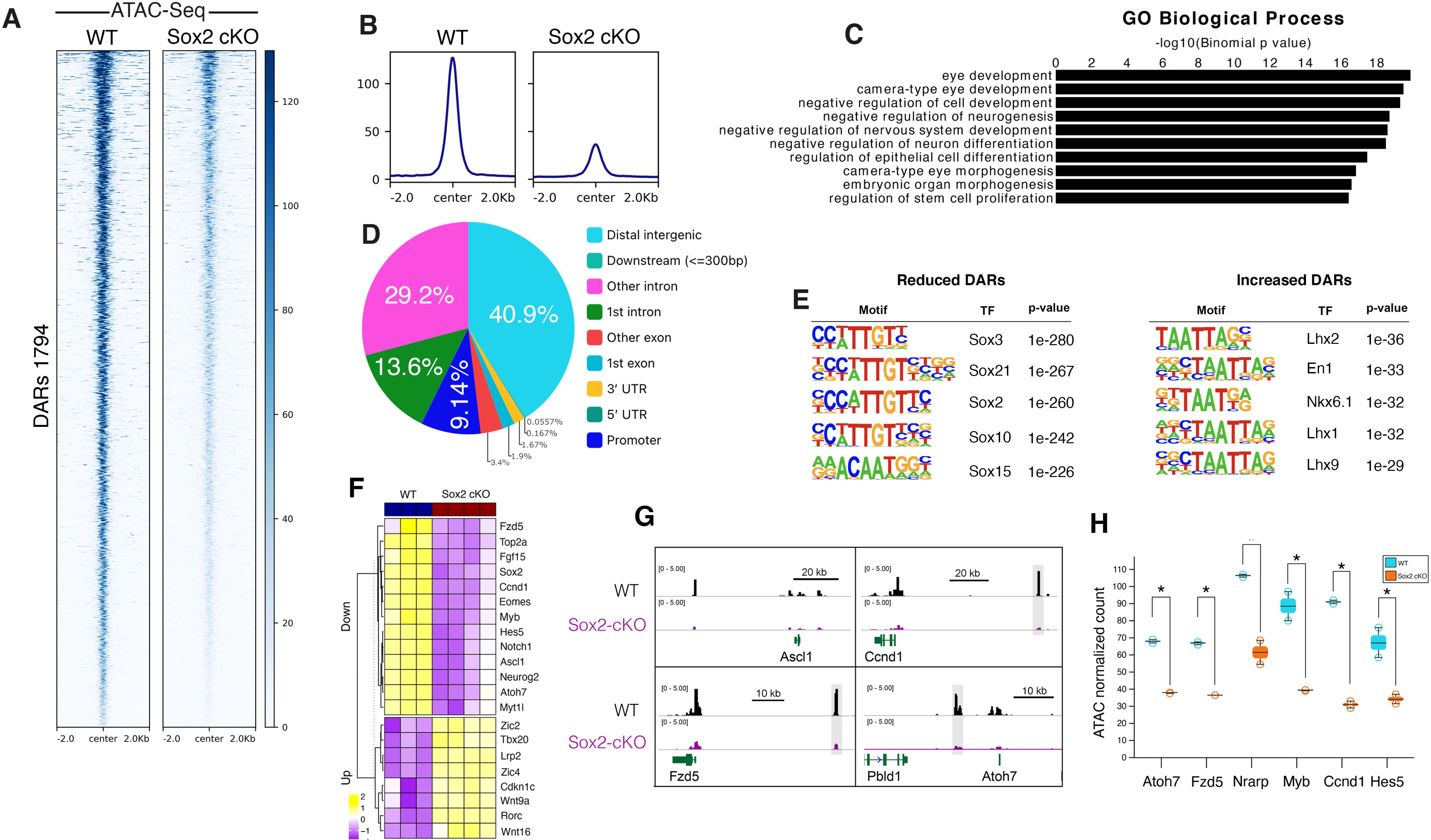
Loss of SOX2 in RPCs disrupts chromatin accessibility. (A) Clustered heatmaps showing differential chromatin accessibility regions (DARs) between WT and *Sox2* cKO retinae at E14.5 (only reduced peaks are shown). (B) Average ATAC-Seq read density profiles in WT and *Sox2* cKO retinae at E14.5. (C) GO biological process analysis of genes associated with reduced DARs in *Sox2* cKO retinae. (D) Pie chart showing the genomic distribution of reduced DARs in *Sox2* cKO retinae. (E) Motif analysis of reduced and increased DARs from *Sox2* cKO. The enrichment of top 5 binding motifs per sample is shown. (F) Heat map showing examples of deregulated genes in Sox2 cKO. (G) Genome browser tracks showing SOX2 binding in WT and ATAC peaks from WT and *Sox2* cKO retinae at key proliferation and differentiation genes. Quantification of the highlighted enhancer regions is shown in H.

To examine whether changes in chromatin accessibility upon loss of SOX2 are concordant with gene expression, we first investigated global changes in gene expression upon loss of *Sox2* in RPCs. We performed RNA-seq on E14.5 retinae from WT and *Sox2* cKO embryos and defined 2180 and 2351 genes that were significantly upregulated and downregulated, respectively (n=3 for WT, n=4 for *Sox2* cKO, adjusted p-value <0.05, > 1.5-fold change) (Fig. S1). *Sox2* cKO developing retinae displayed a loss of proliferation and neural potential as gene networks underlying cell cycle (i.e., *Myb*, *Birc5*, *Foxm1,* and *Ccnd1*) and differentiation (i. e., *Ascl1*, *Neurog2,* and *Atoh7*) genes were significantly reduced (Fig. 1F). Functional enrichment analysis revealed that downregulated genes were mainly enriched in neurogenesis and nervous system development functions, whereas upregulated genes were associated with signaling pathways, particularly Wnt components (Fig. S1). We noted that the Wnt signaling receptor *Fzd5* was among the significantly downregulated genes, indicating that it is a downstream target of *Sox2* in the mouse-developing retina (Fig. 1). This contrasts with data from the *Xenopus* retina, where *Fzd5* regulates *Sox2* expression and suggests species-specific differences in the regulatory network underlying retinogenesis[38].

We next integrated our gene expression data with chromatin accessibility profiles. Within a 50 kb window around the transcription start site (TSS) of expressed genes, we identified 283 deregulated genes that are associated with DARs with the majority of them (221/283; 78%) associated with reduced expression and highly enriched in key neurogenesis-related genes such as *Atoh7, Hes5,* and *Ccnd1*, among others (Fig. 1).

### SOX2 targets the active RPC-specific cis-regulome

To investigate how SOX2 genomic occupancy relates to the RPC chromatin accessibilty, we conducted SOX2 Cleavage Under Targets & Release Using Nuclease (CUT & RUN) assay in duplicate on wild type (WT) mouse retina collected at E14.5 (Fig. 2A). Reads were filtered and aligned to the mouse (mm10) reference genome, and a standard pipeline was used to call peaks (Fig. 2A). We identified 3516 peaks with the vast majority (71.5%) residing outside promoter regions, suggesting strong enrichment of putative non-coding regulatory elements (Fig. 2B). Motif scanning of SOX2-bound peaks yielded the SOX consensus binding motif as the most significant overrepresentation sequence, suggesting a direct DNA binding of SOX2 genomic targets (Fig. 2C). GO data indicated robust enrichment in retinal developmental processes such as eye morphogenesis, epithelial cell differentiation, axis specification and regulation of Notch signaling, including *Notch1*, *Hes5*, *Atoh7*, *Ascl1* and *Neurod1*, in line with key roles of Sox2 [11, 30, 34, 39] (Fig. 2D).

**Figure 2.**
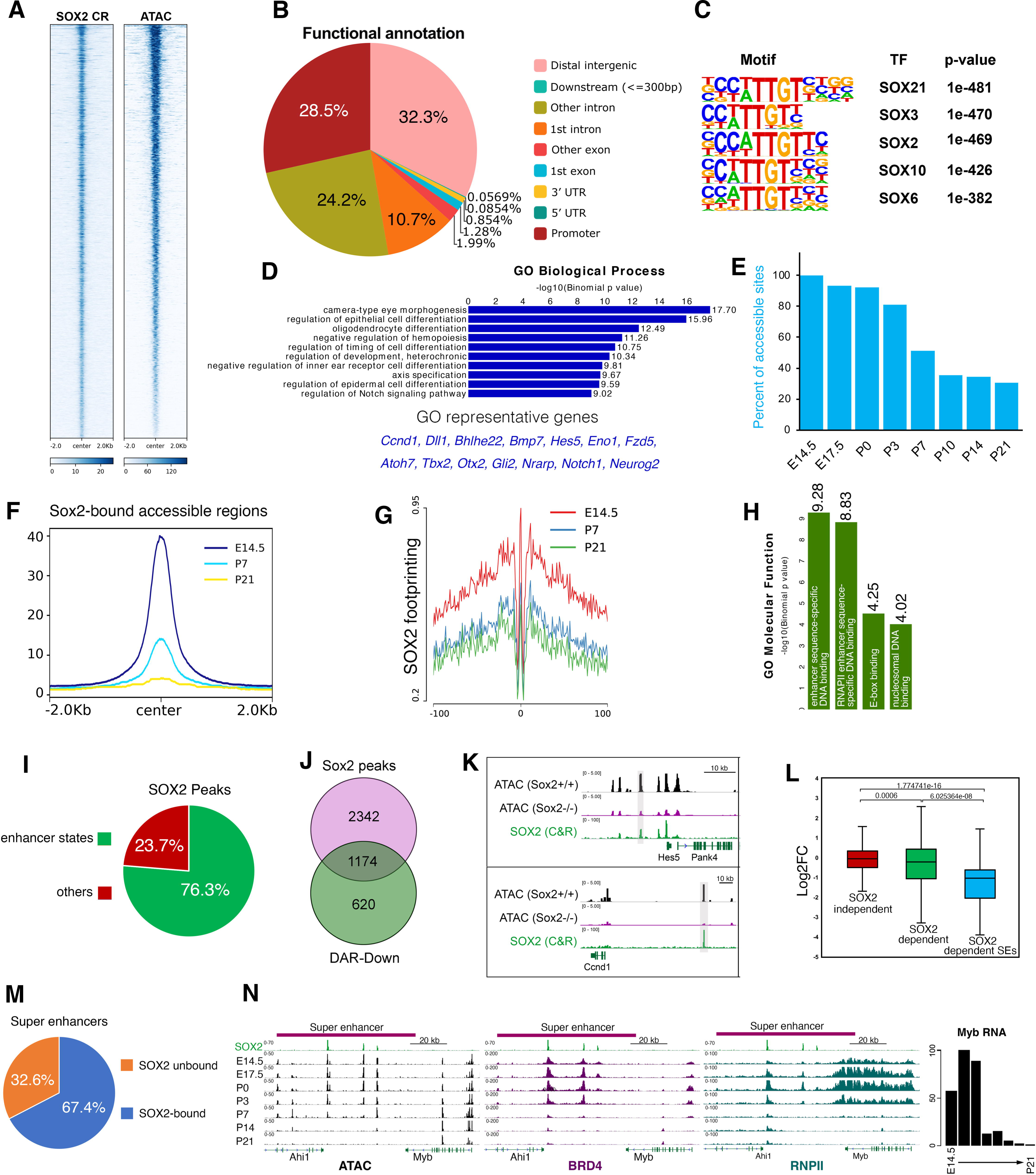
SOX2 targets the active cis-regulatory elements in RPCs. (A) SOX2 CUT&RUN and ATAC-seq binding signal intensity heatmap plotted ±2kb around peak center. (B) Pie chart showing the genomic distribution of SOX2 peaks. (C) Top 5 motif enrichment sequences in SOX2 bound peaks identified from SOX2 CUT&RUN at E14.5 retinae. (D) GO enrichment analysis of SOX2 bound peaks along with representative genes. (E) Bar graph showing the percentage of SOX2-bound accessible sites (SARs) across retinal development (E14.5 to P21). (F) SOX2-bound accessible regions read coverage over three retinal development stages (E14.5, P7, and P21). (G) SOX2 motif footprinting plots at E14.5, P7 and P21 of the developing retina. (H) GO analysis (molecular functions) for SOX2-bound regions. (I) Pie charts showing the proportion of SOX2 peaks associated with enhancer states. (J) Venn diagram showing intersection between SOX2 peaks and reduced DARs in Sox2 cKO. (K) Genome browser tracks showing ATAC signal in WT and Sox2 cKO, SOX2 genomic occupancy. (L) Box plot showing the fold change in gene expression (log2FC) for genes near SOX2-independent, SOX2-dependent, and SOX2-VSX2 co-bound reduced DARs. (M) (left) and the proportion of super-enhancers bound by SOX2 in the developing retina (right). (N) Genome browser tracks showing SOX2, ATAC, BRD4, and RNAPII signals at the *Ahi1-Myb* locus during retinal development. The RNA levels of *Myb* at the corresponding stages are also shown.

We then overlayed SOX2 occupancy with open chromatin regions in WT E14.5 retina and found that 97.6% (3435/3521) of SOX2-bound sites were within accessible sites, revealing a strong correlation between SOX2 binding and chromatin accessibility during the period of high Sox2 expression (Fig. 2A). To examine whether SOX2-bound accessible regions (termed SARs hereafter) are developmentally regulated, we surveyed the dynamics of the SOX2-occupied accessible regions across retinal development using our ATAC data from eight stages (E14.5-P21) [35]. We found that SARs exhibited substantial remodeling upon differentiation and by stage P21 (fully differentiated retina), 69.3% (2381/3435) of those peaks became decommissioned (Fig. 2E-F). To complement this analysis, we investigated SOX2 motif footprint kinetics across retinal development using HMM-based Identification of Transcription factor footprints ATAC (HINT-ATAC)[40]. We found that SOX2 binding exhibited the strongest footprint at E14.5 and gradually decreased afterward, consistent with the dynamics of SOX2-associated accessible regions (Fig. 2G).

GO analysis based on molecular function suggests that SOX2 genomic occupancy was primarily associated with functions related to enhancer activation and decommissioning (Fig. 2H). To investigate this in-depth, we profiled the chromatin status of the SOX2-bound regions using our annotated ChromHMM enhancer states of the developing retina[35]. The analysis denoted that 76.3 % (2682/3515) of SOX2 sites in RPCs were characterized by the acquisition of active enhancer states, indicating that SOX2 is predominantly associated with active cis-regulatory elements in RPCs (Fig. 2I). Importantly, the SOX2-bound enhancer regions carry a strong temporal-specific identity as by stage P21, 92.2% (2421/2628) of those loci exhibit diminished activities.

Intersecting SOX2 peaks with DARs from *Sox2* cKO retina indicated that SOX2 is exclusively associated with reduced accessibility sites as 1174/1794 (65%) of those were bound by SOX2 (named SOX2-dependent DARs hereafter) (Fig. 2J-K). Those sites are enriched in cell proliferation and differentiation genes, suggesting direct role of SOX2 in activating those genes. Indeed, when we compared the log2 fold change (log2FC) of the genes annotated to SOX2-dependent DARs (n = 1174 peaks) to those that are associated with the remaining SOX2 peaks (n = 2342 peak), we found that genes in the neighborhood of SOX2-dependent DARs showed a significant reduction in gene expression over those that are proximal to the remaining peaks (SOX2-independent), thus establishing a direct functional link between SOX2 occupancy, chromatin accessibility and gene regulation (Wilcox test, p-value < 0.05; Figs. 2L).

Among regulatory elements, super-enhancers (SEs) represent a sub-class that is characterized by a very high enrichment of transcription factors and are associated with the expression of key developmental and cell identity genes[41, 42]. We explored whether SOX2 preferentially targets this class of regulatory elements and found that 67.1% (211/314) of super-enhancers in the developing retina were enriched with SOX2 (Fig. 2M-N). One example of those enhancers is located at the *Ahi1*-*Myb* locus (Fig 2N). *Myb* is robustly expressed in the developing retina in RPCs and is downregulated upon loss of *Sox2* (Fig. 1F, 2N). This enhancer region is annotated as a super-enhancer, bound by SOX2 at multiple sites and showing high ATAC, H3K27ac, RNA polII and BRD4 signals that are well correlated with *Myb* expression (Fig. 1N) [43]. We then tested whether SOX2 is an essential regulator of retinal SE-associated transcriptional activities. We charted SEs that exhibited SOX2-dependent reduction in chromatin accessibility and found that 115/211 (54.5%) of those sites were significantly affected. Using gene annotation and manual curation we identified 58 genes that were associated with those SOX2-dependent SEs (Dataset S4), many of which represent integral members of the transcription network that underlines retinogenesis, including *Sox8*, *Prex1*, *Ascl1*, *Myb*, *Fzd5*, *Cxcr4*, *Foxn4*, *Notch1* and *Ccnd1* (Fig. 2L-M). Importantly, those genes showed a strong significant reduction in gene expression when compared to the remaining SOX2-dependent genes, indicating that SOX2-associated SE genes are particularly sensitive to loss of *Sox2* in the developing retina (Wilcox test, p-value<0.05; Fig. 2L). Altogether, these results indicate an essential role of SOX2 in activating the RPC chromatin landscape, as loss of SOX2 leads to a selective disruption of chromatin accessibility and enhancer activities necessary for retinogenesis.

### Putative SOX2-bound cis-regulatory regions are active in the developing retina

While charting the genomic landscape of histone marks and profiling accessibility is a valuable tool to identify chromatin features and pinpoint regions with potential regulatory roles, it falls short in determining whether these regulatory elements are actually functional [44, 45]. Thus, we tested a cohort of SOX2-occupied putative regulatory regions to determine whether they represent *bona fide* enhancers by employing an *ex vivo* reporter assay (Fig. 3) [46–49]. Putative enhancers were selected based on SOX2 occupancy, enrichment of the active enhancer mark H3K27Ac, ATAC-seq signal for chromatin accessibility, and their genomic proximity to key genes involved in retinogenesis. Here, enhancer segments were cloned into Stagia3 vector, which harbors a bicistronic reporter gene cassette (GFP-IRES-PLAP) under the control of a minimal promoter [47]. The vector is *ex vivo* electroporated into dissected WT E14.5 retinae along with a CAG-mCherry plasmid (a constitutively expressing mCherry reporter) as a control for the electroporation efficiency. The electroporated retinae were cultured for 2 days and successful transfectants were screened for mCherry expression and assayed for reporter activity (alkaline phosphatase (AP); Fig. 3). Electroporation of empty Stagia3 vector displayed no AP staining in the retina and serves as a negative control (Fig. 3). Using this strategy, we validated 9 regulatory elements nearby genes that are members of the regulatory network that is associated with retinal proliferation and differentiation, including, *Nrarp (Notch component)*, *Ano1*, *Mybl1*, *Ascl1*, *Ccnd1, Smarcc1* and *Neurog2* (Fig. 3). Overall, enhancer functional validation assay results are consistent with the SOX2 genomic occupancy on putative active RPC enhancers and underscoring the regulatory potential of SOX2 in controlling enhancer-driven gene expression, reinforcing its role as a master regulator of retinal progenitor cell differentiation and identity.

**Figure 3.**
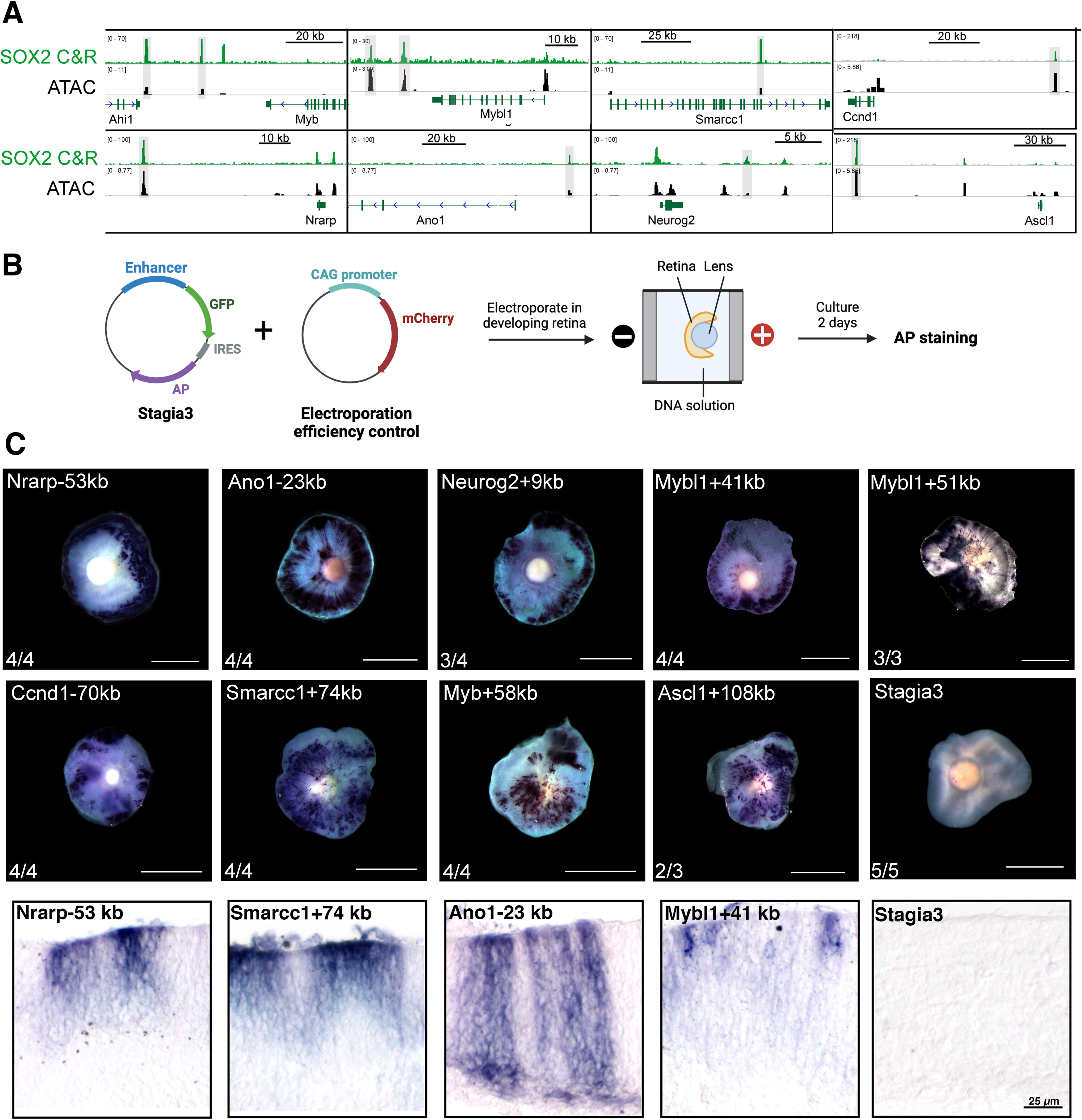
Functional validation of SOX2-bound enhancers using ex vivo reporter assay. A) Genome browser tracks of SOX2 CUT&RUN and ATAC-seq at various loci in the developing retina. (A) (B) Illustration depicting procedure for Stagia3 plasmid electroporation of cultured retinal explant. (B) Representative images of AP activity in E14.5+2DIV retinae after electroporation of SOX2-bound enhancer regions (highlighted in A). The number of retinae with positive AP staining for each enhancer is also shown. Scale bar: 1000 μm.

### VSX2 and SOX2 co-regulate the RPC enhancer landscape

VSX2, a homeodomain-containing transcription factor expressed in RPCs and is essential for retinal proliferation and differentiation [17, 50–53]. Leveraging our previously published VSX2 ChIP-seq data from the developing retina [5], we identified multiple instances of shared genomic binding of SOX2 and VSX2, suggesting potential convergence on critical regulatory pathways governing retinogenesis [54, 55]. To investigate this further, we examined the extent to which SOX2 and VSX2 interact at the chromatin level. By intersecting SOX2 CUT&RUN data with VSX2 genomic occupancy [5], we identified 1,078 shared peaks, 96% (1,038/1,078) of which resided within accessible chromatin regions and were marked by active enhancer signatures (Fig. 4A-B). Notably, these SOX2-VSX2 co-occupied peaks were enriched near genes strongly associated with eye development and cell fate commitment, including *Atoh7*, *Ebf3*, and *Hes5* (Fig. 4D). D*e novo* motif scanning revealed homeodomain and SOX recognition sequences as the top two enriched motifs within the shared sites (Fig. 4C). Interestingly, chromatin binding intensity at co-occupied regions was significantly higher than at sites bound exclusively by SOX2 or VSX2 (Wilcoxon test, *p* < 0.05), further supporting genome-wide cooperativity between these transcription factors (Fig. 4F, G). (Fig. EF-G). Additionally, chromatin accessibility at these shared peaks was dynamically regulated, with the majority being progressively lost during later stages of retinogenesis (only 158/1,038 remained accessible by postnatal day 21).

**Figure 4.**
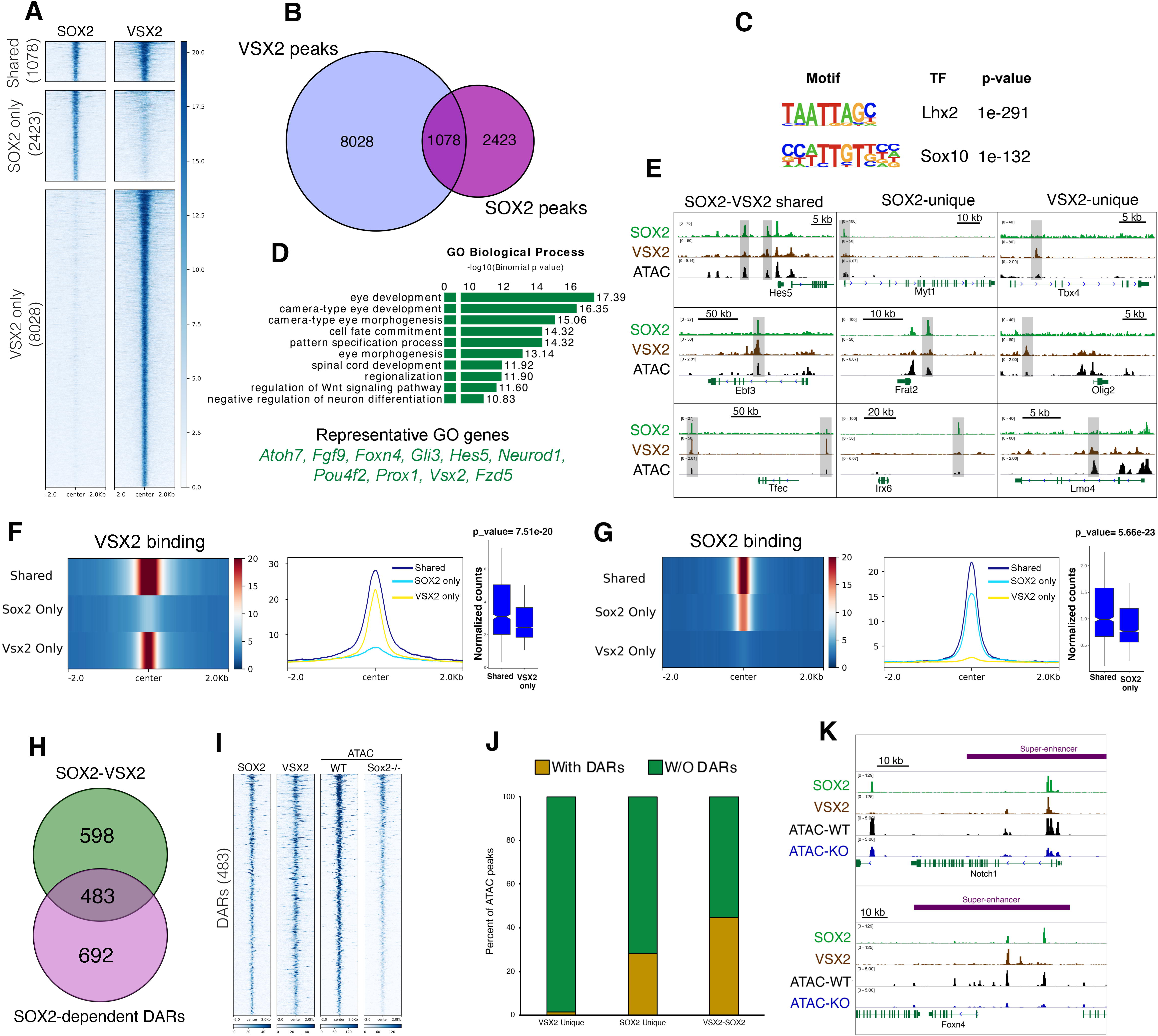
SOX2 and VSX2 co-regulate RPC enhancer landscape. (A) Tornado plots comparing SOX2 and VSX2 genomic occupancy at E14.5 retinae. (B) Venn diagram illustrating the overlap between SOX2 and VSX2 peaks at E14.5 retinae. (C) de novo motif enrichment analysis of SOX2-VSX2 co-bound peaks showing top two hits. (D) GO term analysis of SOX2-VSX2 co-bound associated genes. (E) Genome browser tracks showing examples of SOX2-VSX2 shared, SOX2-unique, and VSX2-unique peaks (highlighted in gray). (F-G) Compressed heatmap (left), average plot (middle) and normalized read counts (right) showing that the strongest binding intensity of VSX2 (F) and SOX2 (G) occurs at shared peaks. (H) Intersection between SOX2-VSX2 shared peaks and SOX2-dependent differentially accessible regions (DARs). (I) Clustered heatmaps comparing SOX2, VSX2, WT and *Sox2* cKO ATAC-seq at reduced DARs. (J) Bar plot showing the percentage of VSX2-unique, SOX2-unique, and SOX2-VSX2 co-bound peaks that occupy SOX2-dependent differentially accessible regions. (K) Genome browser tracks showing SOX2 CUT&RUN and VSX2 ChIP-Seq, ATAC-seq (from WT and *Sox2* cKO retinae) at super-enhancer regions near *Notch1* and *Foxn4*.

We then asked how SOX2-VSX2 genome co-occupancy is related to SOX2-regulated chromatin accessibility. We intersected common SOX2-VSX2 peaks with SOX2-dependent DARs from *Sox2* cKO retina and found that most of the affected peaks (483/1174; 41.1%) co-bound by VSX2 and SOX2, representing 44.7% of total SOX2-VSX2 shared sites (Fig. 4H-J). Within this context, 696 (28.4%) of DARs were characterized by SOX2-unique occupancy, and only 119 (1.5%) were uniquely occupied by VSX2 (Fig. 4J). We also found that 53/211 (25.1%) of SEs occupied by SOX2 engage VSX2, including sites nearby retinogenesis-associated genes such as *Eno1*, *Myb*, *Fzd5*, *Notch1* and *Foxn4* (Fig. 4K).

### SOX2 and VSX2 orchestrate an RPC transcription program and physically interact

We then characterized the underlying regulatory circuits mediated by this SOX2-VSX2 at the transcriptional level in the developing retina, hypothesizing that SOX2 and VSX2 cooperate to regulate a core transcription program in RPCs. We compared differentially expressed genes (DEGs) in E14.5 retinae isolated from the *Sox2* cKO to those previously generated from a *Vsx2* mouse enhancer KO (named Vsx2-EN1 hereafter) at E14.5 retinae, which harbors a homozygous deletion in a regulatory element required for *Vsx2* expression[5]. Vsx2-EN1 KO retinae exhibit a significant loss in proliferation and differentiation potential analogous to *Sox2* cKO mice[5, 56, 57]. We found a substantial overlap between the DEGs in the two lines (Figs. 5A-F). Specifically, 1077 (45.8%) and 971 (44.5%) of downregulated and upregulated transcripts (in *Sox2* cKO), respectively, were shared between *Sox2* cKO and Vsx2-EN1 KO retinae (Fig. 5A-B). GO analysis indicated that shared downregulated genes were heavily linked to nervous system development and neurogenesis functions whereas shared upregulated genes were associated with regulation of cell communication and signaling pathways (Fig. 5C-D). Indeed, signature genes of RPC proliferation and differentiation were enriched within *Sox2*- and *Vsx2*-downregulated genes and included *Notch*, *Ebf3*, *Ascl1*, *Ccnd1* and *Hes5* (Fig. 5E). We also observed significant upregulation of genes normally enriched in the ciliary marginal zone (CMZ) such as *Otx1* and *Gja1*, and Wnt and Bmp signaling pathway components (i.e., *Wnt9a* and *Bmp7*) (Fig. 5H). Interestingly, *Sox2*-*Vsx2* commonly regulated genes were more significantly affected in the *Sox2* knockout than those unique to loss of *Sox2* or *Vsx2* (Wilcox test, p_value<0.05; Fig. 5G-H).

**Figure 5.**
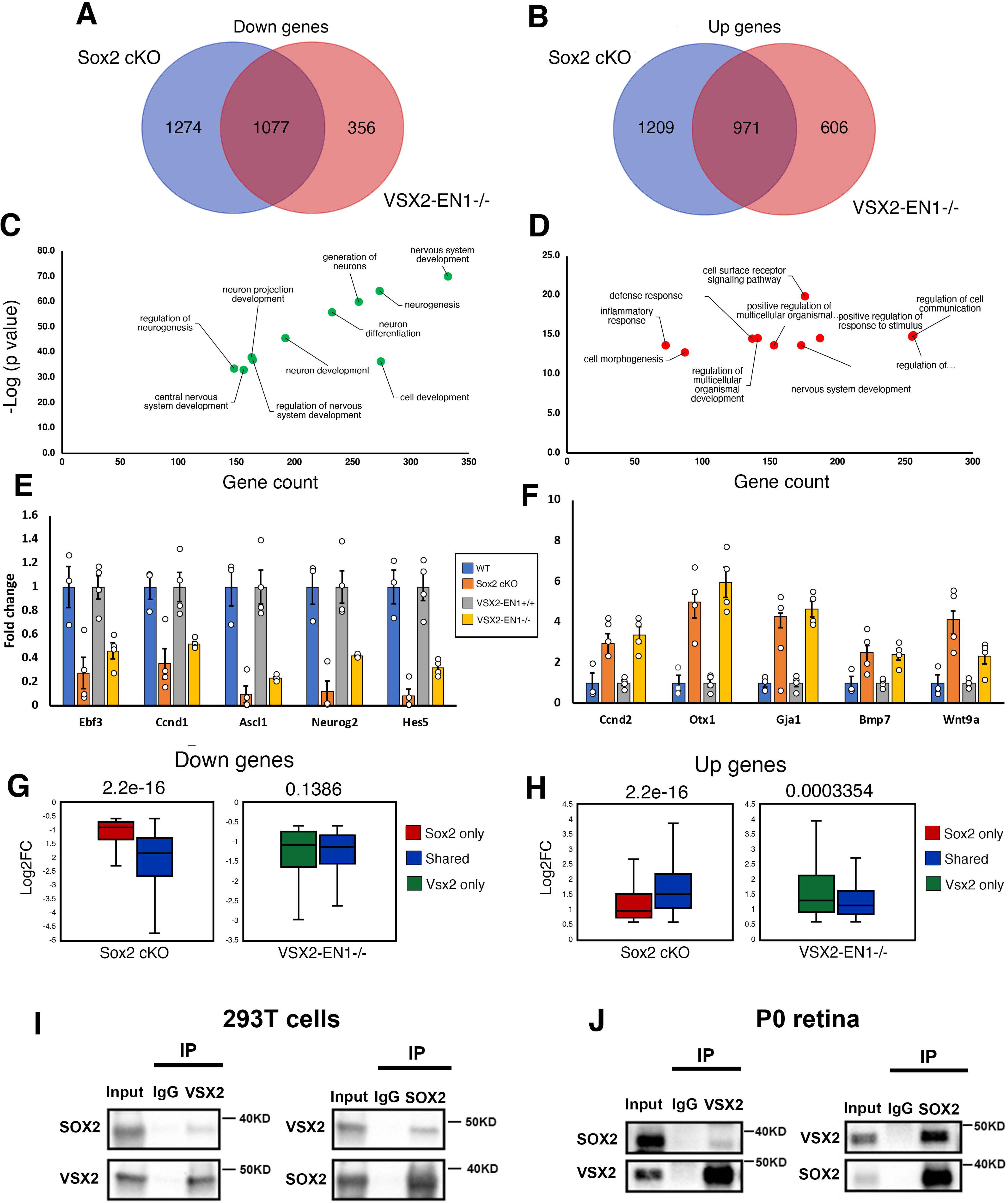
SOX2 interacts with VSX2 to co-regulate RPC transcriptional programs. (A-B) Venn diagrams showing the overlap between down- (C) and up-regulated (D) genes in *Sox2* cKO and Vsx2-EN1 KO retinae. (C-D) GO enrichment analyses of shared down- (E) and up-regulated (F) genes in *Sox*2 cKO and Vsx2-EN1-/- retinae. (E-F) Bar plots showing fold changes in gene expression for representative common down and up-regulated genes in *Sox2* cKO and Vsx2-EN1 KO retinae. (G-H) Box plots comparing the log2 fold changes (Log2FC) of SOX2-VSX2 shared versus *Sox2*- and *Vsx2*-unique transcriptional targets for downregulated (I) and upregulated (J) genes (Wilcox test, p-value <0.05). (I-J) Co-IP assays on cell lysates of SOX2 and VSX2 transfected 293T cells (I) and P0 retinae (J) with relevant quantifications. 3% of input was used.

Based on these data we tested whether SOX2 and VSX2 physically interact. We first performed immunoprecipitation assays using lysates from 293T cells that co-express VSX2 and SOX2 and found that SOX2 efficiently co-immunoprecipitated with VSX2 (Fig. 5A). To confirm this interaction *in vivo* where SOX2 and VSX2 are expressed at physiological levels, we performed reciprocal immunoprecipitation assays on the developing retina at P0 and were able to validate the physical interaction of both transcription factors (Fig. 5B). Together, our data indicate a mechanism by which the SOX2-mediated transcriptional program during retinogenesis involves cooperation with VSX2 to promote proliferation and subsequent differentiation.

### SOX2 and VSX2 co-occupy *Vsx2* regulatory elements

We investigated the *Vsx2* regulatory landscape which contains a modular enhancer with activity that is associated with *Vsx2* expression *in vivo*[5, 57]. *Vsx2* enhancer structure contains two conserved hot spots, termed EN1 and EN2, that are required non-redundantly for retinal development and *Vsx2* expression (Fig. 6A)[5, 57]. VSX2 occupies and regulates the activity of EN1 and EN2, underlying an autoregulatory loop that governs *Vsx2* expression (Fig. 6A)[5]. Here, we discovered that SOX2 also robustly binds EN1 and EN2, suggesting that regulation of the *Vsx2* enhancer landscape may also involve SOX2 (Fig. 6A). To rigorously investigate EN1 and EN2 activity *in vivo* we generated transient transgenic mice harboring EN1 or EN2 elements using enSERT, an innovative mouse transgenic assay that allows robust testing of enhancer activities in mice with high efficiency[58]. Consistent with our data from *ex vivo* electroporation assays, E12.5 embryos harboring EN1 or EN2 transgenes showed robust LacZ staining in the developing retina (Fig. 6A; 5/5 embryos for EN1 and 4/5 embryos for EN2).

**Figure 6.**
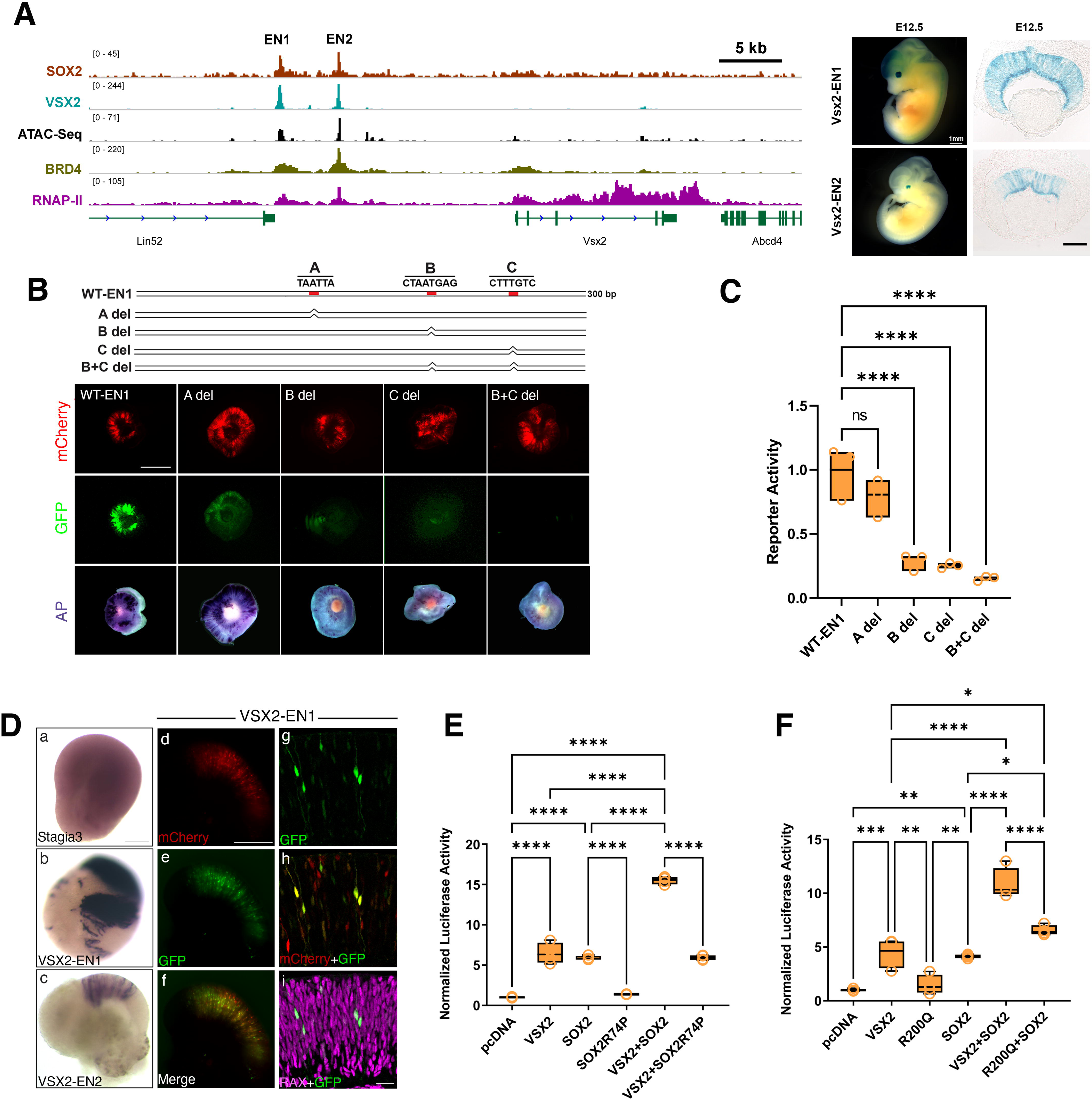
SOX2 and VSX2 co-regulate Vsx2 enhancer activity in RPCs. (A) Chromatin landscape near *Vsx2* with SOX2, VSX2, BRD4, RNAP II and ATAC signals at E14.5 (right). enSERT transgenic mice harboring the *Vsx2* regulatory elements EN1 and EN2 (left). LacZ staining of embryos (Scale Bar: 1mm) and corresponding eye sections at E12.5 (Scale Bar: 100 μm) are shown. (B) Top panel: Schematic of the Vsx2-EN1 enhancer showing the relative position of potential transcription factor binding motifs A, B, and C. Lower panel: GFP signal and AP staining results alongside mCherry efficiency for WT-EN1 (300bp), A motif deletion (A del), B motif deletion (B del), C motif deletion (C del), and B+C double motif deletion (B+C del). Image taken at E14.5+2 DIV. Scale bar: 500 μm. (C) Quantification of mutant VSX2-EN1 enhancer activities. The ratio of total GFP intensity over mCherry intensity was calculated and normalized to VSX2-EN1. ****P<0.0001. (D) AP staining of Day 44+ 2 DIV Human retinal organoids differentiated from a control line and electroporated with empty Stagia3 plasmid (a) or one containing Vsx2-EN1 (b) or Vsx2-EN2 (c) (Scale bar 200 μm). mCherry (d) and GFP (e) signals from hRetOrg co-electroporated with Stagia3 plasmid containing Vsx2-EN1 along with mCherry (scale bar 150 μm). (g-h) Confocal images of D44 + 2 DIV human retinal organoids showing overlap between GFP, mCherry and the RPC marker RAX (scale bar 30 μm). (E) Luciferase assay of human Vsx2-EN1 in the presence of VSX2, SOX2R74P, SOX2, VSX2+SOX2, VSX2+ SOX2R74P. Data are mean±s.e.m., n=4. ****P<0.0001 ***P<0.001; **P<0.01, *P<0.05, one-way ANOVA. (F) Luciferase assay of human Vsx2-EN1 in the presence of SOX2, VSX2, VSX2 R200Q, VSX2+SOX2, VSX2 R200Q+ SOX2. Data are mean±s.e.m., n=4. ****P<0.0001 ***P<0.001; **P<0.01, *P<0.05, one-way ANOVA.

### Analysis of SOX2 and VSX2 motifs on *Vsx2* enhancer

We then used a bioinformatics approach to identify candidate sequences in the *Vsx2* enhancer landscape that might confer RPC enhancer activity, hypothesizing that SOX2 and VSX2 directly bind EN1 and EN2 via discrete sequences. To simplify the analysis, we focused on EN1, as our group and others have demonstrated that it is essential for RPC proliferation in mice[5, 57]. Here, we narrowed our search to a 300 bp region in EN1 that successfully activated AP reporter expression in E14.5 retina (WT-EN1; Fig. 6B; Fig. S5). Using the catalog of inferred sequence binding preferences (Cis-BP) database, we identified three potential TF motifs, two of which were related to homeodomain-containing binding sequences (possible VSX2 binding sites; termed hereafter A and B), and a third sequence that contained a SOX-related binding site recognition motif CTTGTC, (termed C hereafter; Fig. 6B). To functionally examine the significance of these motifs, we deleted each of them individually and tested the mutant constructs with our retinal electroporation assay. We found that whereas the A motif was dispensable for AP activities (4/4 retina with positive AP staining) both B and C had a significant reduction in AP staining (4/4 retina showed a reduction in AP; Fig. 6B). We also generated a compound mutant construct in which both B and C motifs were eliminated, resulting in a further reduction in activities (4/4 retina showed a reduction in AP, Fig. 6B). In summary, our analysis implicates SOX2 and VSX2 in the regulation of the Vsx2 enhancer landscape, underlying complex combinatorial interactions among transcription regulatory proteins in driving RPC enhancer activation.

### VSX2 super-enhancer activities are conserved in human retinal organoids

The VSX2 enhancer sequence is conserved in humans but whether this element is sufficient to drive activities in the developing human retina is unknown. Retinal organoids generated from human stem cells offer an unprecedented opportunity for studying human retina development, identifying mechanisms of disease, and facilitating the discovery of new treatments[59–61]. Importantly, immunohistochemical and single-cell transcriptomics of the developed organoids have confirmed their high similarity to the human fetal retina and recapitulation of many *in vivo* developmental programs, providing a foundation for understanding human retina development and disease *in vitro*[61]. Taking advantage of this system, we developed a scalable protocol to generate human retinal organoids (hRetOrg) from human induced pluripotent stem cells (hiPSCs) and to systematically test the enhancer activities identified in mouse studies using our *ex vivo* electroporation method. hRetOrg from a commercially available control hiPSC line (IMR90.4) were grown to day (D) 44-D50, corresponding to early stages of retinogenesis (∼8-week-old human fetal retina). We then electroporated individual Vsx2-EN1, -EN2 or empty Stagia3 plasmids with pUbC:mCherry vector and cultured them for two days (Fig. 6D, Fig. S5). We then subjected the electroporated organoids to AP staining and found robust AP expression in both EN1 and EN2 (n=10 organoids with positive signal for each construct; 0/10 for empty Stagia3 organoids). Indeed, GFP signal indicated that the VSX2-EN1 was selectively activated in cells with morphology similar to RPCs, and co-localized with RAX, a canonical marker of RPCs (Fig. 6D). Our data established hRetOrg as a highly scalable and reliable platform to test enhancer activities and indicates that the mouse VSX2-EN1 element is functionally conserved in humans.

### Co-activation of *Vsx2* enhancer elements by SOX2 and VSX2

To test whether SOX2 can influence the activity of the VSX2-EN1 element we performed luciferase reporter assays on 293T cells. We used the human *VSX2-EN1* sequence in this analysis as it is more clinically relevant. We previously demonstrated that VSX2 significantly stimulated Vsx2-EN1 luciferase activity, an effect that was virtually eliminated when a VSX2 mutant that is associated with microphthalmia (R200Q) was utilized[5]. Here, we discovered that SOX2 also significantly induced EN1 activity (n=4, ANOVA test, P<0.0001) (Fig. 6D, E). Co-expression of VSX2 and SOX2 potentiated EN1 transcriptional activity, well above the additive effect of combined SOX2 and VSX2, indicating that the two classes of TFs can synergize in regulating gene expression (n=4, ANOVA test, P<0.0001; Fig. 6D, E). The introduction of an AM-associated SOX2 mutation (R74P), which harbors a single amino acid substitution in the DNA-binding domain (HMG)[62], abolished SOX2’s ability to activate EN1 and impacted the synergistic effect between VSX2 and SOX2 (n=4, ANOVA test, P<0.0001; Fig. 6D, E). These data further indicate that SOX2 is involved in the regulation of *Vsx2* expression in coordination with VSX2.

## Discussion

RPC differentiation potential is driven, at least in part, by the highly specific regulatory interplay between RPC cis-regulatory modules and underlying gene networks; however, the molecular underpinnings of this process remain enigmatic. Here, we focused on deciphering how SOX2, a transcription factor (TF) required for RPC proliferation and differentiation and a core member of the RPC gene regulatory network, functions at the genome-wide level to establish the RPC chromatin landscape. We found that SOX2 is highly enriched in nucleosome-depleted regions distal to promoters of key genes essential for retinal proliferation and differentiation. These sites are characterized by the acquisition of active enhancer marks and progressively become inactive as retinal differentiation proceeds, revealing that SOX2 primarily targets developmentally regulated enhancers underlying the RPC neural identity.

Evidence indicates that SOX2-mediated control of retinal proliferation and differentiation can be uncoupled and involves distinct molecular mechanisms [34, 63]. We demonstrate that SOX2 activation of retinogenesis entails direct regulation of chromatin accessibility across the enhancer landscape of crucial developmental genes, such as proneural bHLH factors and components of Notch signaling. Mechanistically, SOX2 modulation of chromatin accessibility in the retina likely requires the recruitment of chromatin remodeling complexes that reposition nucleosomes. Indeed, evidence from other systems demonstrates that SOX2 interacts with subunits of the SWI/SNF complex [28, 64]. However, whether SOX2-mediated neural competency directly depends on chromatin remodeling activity remains unclear. For instance, deletion of the SWI/SNF core subunit *Samrca4/Brg1*, which is expressed in the developing retina, leads to proliferation defects, though cell type specification proceeds normally [65]. Furthermore, chromatin remodeling factors that interact with SOX2 in RPCs to facilitate accessibility have not yet been identified.

Our findings underscore a pivotal role for SOX2 in governing chromatin accessibility. SOX2 may be required to initiate accessibility by unlocking compacted chromatin regions, potentially through direct engagement with nucleosomal DNA and recruitment of chromatin remodelers and histone-modifying enzymes. Alternatively, SOX2 may stabilize pre-existing open regions, acting in concert with transcription factors and epigenetic regulators to sustain an open chromatin state and ensure the fidelity of gene expression programs. These functions are not mutually exclusive, as SOX2 could contribute to both processes. Notably, SOX2’s interaction with chromatin remodeling complexes is well documented, and our data reveal its binding to the transcription factor VSX2, a key regulator of retinogenesis. However, since our analysis begins at E14.5, it remains unclear whether SOX2 primarily functions as a pioneer factor, a maintenance factor, or both.

Transcriptionally, SOX2 can sustain RPC proliferation by directly promoting the expression of factors required for cell cycle progression. Supporting this hypothesis, we observed that SOX2 occupies distal regulatory elements near proliferation genes such as *Myb* and *Ccnd1*, and experimentally confirmed that the associated putative enhancers are functional in the developing retina (Fig. 3). Furthermore, loss of SOX2 leads to a significant downregulation of proliferation genes and diminished chromatin accessibility at the corresponding regulatory elements (Fig. 2). Interestingly, contrary to its role during early retinogenesis, SOX2 acts to repress *Ccnd1* expression in a Wnt-dependent manner in the developing cortex, underscoring that SOX2-mediated regulation of *Ccnd1*, and likely other cell cycle regulators, is context-specific [66, 67]. SOX2 may also promote proliferation by repressing cell cycle inhibitors. We discovered that in the *Sox2*-deficient developing retina, there is significant upregulation of the cell cycle inhibitor *Cdkn1c* (2.6-fold increase, adj p-value < 0.05; Dataset S2; Fig. S1), which impedes mitotic RPCs [68]. Altogether, we conclude that SOX2 control of retinal proliferation involves both activation and repression mechanisms.

One of the least understood roles of SOX2 is its capacity to facilitate transcriptional repression, likely through the recruitment of co-repressors [69–72]. Evidence from various tissues suggests that the Wnt pathway is a primary target for SOX2-mediated antagonism[34, 73–75]. In the developing mouse retina, Wnt signaling is normally active in the CMZ, where it promotes the formation of anterior eyecup structures at the expense of the neural retina [34, 76, 77]. Our data supports work indicating that SOX2 directs retinal neural differentiation in part by blocking the expression of Wnt factors [34]. Not all Wnt components, however, are targets for SOX2 repression. For example, the Wnt receptor *Fzd5*, which is expressed in RPCs, is positively regulated by SOX2, likely through direct binding to its enhancer elements (Fig. S1). Our data place *Fzd5* as a downstream target of SOX2, contrasting with data from *Xenopus*, underlining species-specific differences in the transcriptional regulation of RPC differentiation [38].

SOX2 is expressed in embryonic stem cells and many developing tissues, yet its genomic occupancy is highly context-dependent [67]. Evidence suggests that SOX2 target selectivity is achieved through cooperative binding with tissue-specific transcription factors [67, 78]. We uncovered a novel regulatory circuit involving SOX2 and the RPC transcription factor VSX2 that functions on the chromatin level and converges on a common RPC gene architecture essential for retinal neural competency. We postulate that the SOX2-VSX2 partnership constitutes an integral part of the RPC gene regulatory networks that promote neural competence by facilitating SOX2 target selectivity and potentiating transcriptional regulation of key retinogenesis genes. Additionally, our data suggest that the molecular mechanisms underlying SOX2-VSX2 cooperativity also involve transcriptional repression, which appears independent of modulation of chromatin accessibility. The exact role of the SOX2-VSX2 axis in the developing retina, particularly in facilitating gene repression at the chromatin level, remains unclear.

While our data suggest that Sox2 contributes to the activation of part of the Vsx2-EN1 enhancer, we observed that Vsx2 expression was not downregulated upon the loss of Sox2. Although the exact mechanisms are unclear, this suggests the involvement of redundant regulatory mechanisms, potentially including both cis- and trans-acting elements. One possibility is that other transcription factors compensate for the absence of Sox2, maintaining Vsx2 expression. Alternatively, the Vsx2 regulatory landscape is complex, comprising multiple enhancers and regulatory elements. Additional enhancers with binding sites for different transcription factors may help sustain Vsx2 expression even without Sox2. Notably, these mechanisms may not be mutually exclusive and could function in parallel. Overall, our findings highlight the complexity of transcriptional regulation, which often relies on multiple layers of control to ensure robust gene expression.

SOX2 expression persists postnatally in Müller glia (MG), a non-neural cell type with neuroprotective, structural, and metabolic functions essential for maintaining viability, preserving retinal homeostasis, and responding to stress conditions [79, 80]. In mice, *Sox2*-deficient MG re-enter the cell cycle but do not differentiate into retinal cell types, eventually leading to retinal degeneration [30, 81]. We speculate that SOX2 binding partners and genomic targets in MG differ from those in RPCs, involving unique transcriptional regulation of proliferation and stress-response genes, though further studies are required to test this hypothesis.

In summary, our data demonstrate how the regulatory interaction between SOX2 and tissue-specific partners establishes and reinforces the RPC chromatin landscape necessary for retinal cell-type-specific transcriptional programs, providing a molecular mechanism underlying the RPC developmental potential and early cell fate acquisition. **Materials and Methods**

### Animals

#### Mice

All mice were maintained in accordance with the guidelines set forth by the Institutional Animal Care and Use Committee of the University of Pittsburgh. C57 BL/6J, *Sox2^tm1.1Lan^*/J (013093), Chx10-EGFP/cre,-ALPP)2Clc/J (005105) animals, were purchased from Jackson Laboratory. *Sox2* conditional knockout is generated by crossing Sox2 floxed mice^□^with *Chx10* ^Cre-GFP^ mice.

### CUT&RUN

Freshly dissected E14.5 retinae were lysed for 10 minutes at RT in lysis buffer. Nuclei were then bound to Bio-Mag Plus Concanavalin A coated beads (Cell Signaling #86057-3). After blocking with 2mM EDTA blocking solution for 10 mins at RT, beads were washed and then bound to SOX2 antibody (Active Motif #39844) overnight. We used no-primary-antibody as a negative control[82–84]. The next day, samples were treated with pA-MNase and subjected to digestion with the addition of CaCl2. The digestion reaction was stopped after 30 minutes then samples were treated with RNase for 20 mins at 37°C to release CUT&RUN fragments from beads. Fragments were then purified using phenol-chloroform extraction followed by ethanol precipitation. Library preparation was then performed with NEBNext Ultra II DNA Library Kits for Illumina (NEB #E7645, E7103) using 10ng of CUT&RUN fragments. Libraries were multiplexed and run on a high output 75 cycles flowcell with a read length 2 x43, using the sequencing platform NextSeq500.

### CUT&RUN Analysis

Reads were processed using the nf-core-cutandrun pipeline (v3.1)[85]. Quality control of paired end FASTQ files was performed using FastQC. Adapter sequences were removed with Trim Galore (Babraham Institute). Reads were aligned with Bowtie2[86] to mm10 and duplicates were removed for control samples only using Picard (Broad Insitute). Peaks were called using MACS2[87]. Consensus peaks were merged across all samples using BEDTools[88] and annotated to the nearest gene transcriptional start site using HOMER[37]. Peak counts were generated using featureCounts[89].

Analysis performed using platform (https://pluto.bio).

### ATAC-seq preparation and Library Sequencing

ATAC was performed using Active Motif ATAC-seq kit (Catalog No. 53150) according to the manufacturer’s protocol. Briefly, dissociated E14.5 retinal cells were subjected to tagmentation reaction at 37°C for 30 min. DNA purification was carried out with a Qiagen MinElute PCR Purification Kit (Qiagen, 28006) and PCR amplification was performed using indexed primers.The resulting library was quantified and assessed through a Qubit FLEX fluorometer and an Agilent TapeStation 4150 respectively.

Libraries were pooled and normalized to 2 nM. sequencing was carried out on an Illumina NextSeq 2000 using a P1 100 flow cell. With a target of 25 million reads per sample, the pooled library was loaded at 800 pM and sequencing was carried out with read lengths of 2x58 bp. Sequencing data was demultiplexed with the on-board Illumina DRAGEN FASTQ Generation software.

### ATAC-seq Analysis

Reads were processed using the nf-core-atacseq pipeline (v2.0)[85]. Adapter sequences were removed with Trim Galore (Babraham Institute). Reads were aligned using Bowtie2[86] to the reference genome mm10. Following alignment, duplicate reads were removed using the Picard. Peaks representing accessible chromatin regions were called using MACS2[87] (peak mode: narrow) by means of a p-value threshold of 0.00001 and an effective genome size of 2,650,000,000. Consensus peaks across all samples merged utilizing BEDTools[88] and annotated to the nearest gene transcriptional start site using HOMER. Peak counts were generated through featureCounts[89]. Differential accessibility peaks were identified by DEseq2[90].

Analyses were performed using Pluto platform (https://pluto.bio).

### HOMER and GREAT analyses

HOMER enrichment motif analysis was run with standard settings with repeat masking on. 200 bp surrounding the summit of peaks were analyzed. GREAT analysis was done using mm10 (mouse) or hg38 (human) with default parameters.

### RNA extraction and bulk RNA-seq

Fresh retinae were placed in TRIReagent (Zymo Research R2050-1-50), and RNA was extracted according to Direct-zol RNA MiniPrep kit protocol (Zymo Research R2050).

For each sample, 7.5 ng of input RNA was used for library preparation. After adapter ligation, 14 cycles of indexing PCR were performed using Illumina RNA UD Indexes. RNA libraries were normalized and pooled to a final concentration of 2 nM. The Illumina Stranded Total RNA Library Prep kit (Illumina, 20040529) was used following the manufacturer’s protocol for library preparation. Sequencing was conducted on an Illumina NextSeq 2000 using a P3 flow cell. The pooled library was loaded at a concentration of 750 pM, targeting 40 million reads per sample, with read lengths of 2x101 bp. Demultiplexing of the sequencing data was performed using the Illumina DRAGEN FASTQ Generation software.

### Analysis of bulk RNA-seq

Raw sequencing reads first went through the pipeline of quality control [QC] using Trimmomatic. After pre-processing, surviving reads were aligned to mouse reference genome mm10 by STAR aligner and quantified by read count using HTSeq tool.

Differential expression analysis comparing wild-type and knock-out samples were performed by R package DESeq2[90]. Significant changes in gene expression were defined by adjusted p-value lower than 0.05 (FDR=5%) and fold change greater than 1.5. All the bioinformatics and biostatistical analysis were performed by default parameters.

### Luciferase reporter assay

The luciferase reporter assays were carried out following the protocol outlined in our previous publication[5]. In brief, human Sox2 cDNA was subcloned into a modified pcDNA3 vector, where the CMV promoter was replaced with the EF1α promoter. Human Vsx2 cDNA, VSX2R200Q, along with the Vsx2 EN1 enhancer plasmids, were previously described[5]. Mutagenesis of Sox2 was performed using overlap extension PCR, as described previously[91].

HEK293T cells (ATCC® CRL-3216) were maintained in DMEM (Thermo Fisher Scientific, 10569044) supplemented with 10% FBS (Thermo Fisher Scientific, 16000036), 100 U/ml penicillin, and 100 µg/ml streptomycin (Thermo Fisher Scientific, 10378016). Cells were seeded at a density of 30,000 cells per well in 24-well plates and allowed to adhere overnight. For each transfection, 500 ng of either Vsx2, Sox2, Sox2R74P, or R200Q expression constructs were co-transfected with 200 ng of Vsx2 enhancer vectors and 10 ng of the firefly luciferase reporter plasmid pGL4.53[luc2/PGK] (Promega, E5011). For co-expression conditions, 250 ng of Vsx2 or R200Q was combined with 250 ng of Sox2 or R74P and co-transfected with the enhancer and reporter plasmids.

Transfections were performed using Lipofectamine 3000 (Thermo Fisher Scientific, L3000008) following the manufacturer’s instructions. Control transfections included empty vectors co-transfected with each of the enhancer constructs and the firefly luciferase plasmid. Cells were harvested and lysed 24 hours post-transfection, and luciferase activity was measured using the Nano-Glo Dual-Luciferase Reporter (NanoDLR) assay system (Promega, N1610) in accordance with the manufacturer’s protocol. Luminescence was detected using a SpectraMax M5 (Molecular Devices) plate reader. All experiments were performed in quadruplicate.

### Immunoprecipitation

HEK-293T cells were transfected with the respective DNA constructs using Lipofectamine 3000 (Invitrogen) in 6-well plates. Two days post-transfection, cells were lysed in a high-salt lysis buffer (10% glycerol, 50 mM Tris pH 7.5, 300 mM NaCl, 1% NP-40, 2 mM MgCl₂) supplemented with a protease inhibitor cocktail. Lysates were incubated on a rotating platform at 4°C for 30 minutes. To reduce the salt concentration, an equal volume of no-salt lysis buffer (10% glycerol, 50 mM Tris pH 7.5, 1% NP-40, 2 mM MgCl₂) was added to the lysates.

For immunoprecipitation (IP), 500 µg of protein from the lysates was incubated with the appropriate primary antibodies overnight at 4°C. For endogenous IP, non-denatured native protein lysates were prepared from C57BL/6J P0 retinae using COIP buffer (PBS, 10% glycerol, 0.6% Triton X-100) with protease inhibitors. On average, 5 retinae were pooled for each lysate preparation. Equal volumes of the cell or retinal lysates were incubated with the specific antibodies or control IgG overnight at 4°C.

The following day, protein A/G magnetic beads (Thermo Fisher Scientific, 88802) were added to the mixtures and incubated for 3 hours at 4°C. The beads were then washed five times with a low-salt lysis buffer (10% glycerol, 50 mM Tris pH 7.5, 150 mM NaCl, 1% NP-40, 2 mM MgCl₂). Proteins bound to the beads were eluted using SDS-PAGE loading buffer and subjected to SDS-PAGE, followed by transfer to PVDF membranes. Membranes were blocked with 5% non-fat dried milk in TBS-Tween and subsequently incubated with the appropriate primary antibodies. To avoid heavy chain contamination (∼50 kDa), different species of secondary antibodies were used for detection.

### Electroporation and AP staining

Enhancer fragments were amplified from mouse genomic DNA using the primers listed below. These amplified fragments were then cloned into the Stagia3 vector via restriction enzyme-based cloning methods or gibson cloning. Plasmids for electroporation were prepared at a final concentration of 700 ng/µl per plasmid. For electroporation, dissected E15 retinas or Day 44 Human retinal organoids (hRetOrg) were subjected to five pulses at 25V with 50ms on and 950 ms off intervals. Following electroporation, the samples were carefully transferred to 12-well plates and cultured for 2 days. Post-culture, the samples were washed with PBS and fixed in 4% PFA for 30 minutes at room temperature. Before proceeding with alkaline phosphatase (AP) staining, the presence of the mCherry signal was confirmed to assess electroporation efficiency, as fluorescence intensity diminishes significantly after AP staining. GFP signals, if present, were also recorded. The samples were then incubated at 65°C for 1.5 hours, followed by incubation in AP buffer (100 mM Tris-HCl, 100 mM NaCl, 5 mM MgCl₂, pH 9.5) for 15 minutes at room temperature. The buffer was then replaced with developer solution (prepared by dissolving one NBT/BCIP tablet in 10 ml of distilled water). AP signal development was monitored after at least 20 minutes of incubation at room temperature. The reaction was stopped by washing the retinas with PBS, and images of the stained samples were captured using a Leica M165 Stereo Microscope. Quantification of fluorescence images of Fig. 6B was performed using Fiji software by calculating the total fluorescence intensity ratio of EGFP to mCherry, and values were normalized to the WT-EN1 group.

### Transient Transgenic Embryos Reporter Assay (*enSERT*)

The enSERT reporter assay was carried out following previously published methods[58].

Vsx2 enhancer regions EN1 and EN2 were cloned into the Shh::LacZ-H11 vector (Addgene #139098). The targeting vector was then microinjected along with Cas9 protein and a gRNA targeting the H11 locus into the pronuclei of fertilized ova. Embyros were harvested at E12.5 for LacZ staining. Primers used in cloning the VSX2 enhancers are available upon request.

### Human induced pluripotent stem cells (hiPSC) maintenance

The hiPSC line IMR90-4 (WiCell) was utilized for differentiation of human Retinal Organoids (hRetOrg). iPSCs were cultured at 37°C and 5% CO2 in a humidified incubator using mTeSR^TM^ plus medium (STEMCELL Technologies, #100-0276) in 6-well plates (Corning, 3516) coated with hESC-qualified Matrigel (Corning, 354277)and passaged at 70%-80% confluency.

### Differentiation of human Retinal Organoids (hRetOrg)

hRetOrg were differentiated as previously described (Wang etal., 2024) based on the original protocols [Zhong et al., 2014 and Cowan et al. Flickr et al 2020]. Briefly, hiPSC were dissociated to single cells using Accutase (Thermofisher Scientific, # 00-4555-56) and careful mechanical pipetting using a P1000 in 1ml of mTeSR^TM^ plus containing 10µM Rock inhibitor Y-27632 (STEMCELL Technologies, #72304). hiPSC were then plated on 10cm petri dishes (VWR # 25384-088) with a total volume of 10ml of mTeSR^TM^ plus to promote formation of embryoid bodies (EBs). On Day (D) 1, approximately 1/3 of the medium was exchanged for neural induction medium (NIM) containing DMEM/F12 (Gibco, #11330057), 1% N2 supplement (Gibco, #17502048), 1x NEAAs (Sigma, #M7145), and 2mg/ml heparin (Sigma, # H3149). On D2, approximately 1/2 of the medium was exchanged for NIM. On D3 EBs were plated in new 60mm petri dishes (Corning #430166) in NIM. Half of the medium was changed every day until D6. At D6, BMP4 (R&D, #314-BP) was added to the culture at a final concentration of 1.5nM. At D7, EBs were transferred to 60mm dishes previously coated with Growth Factor-Reduced (GFR) Matrigel (Corning, #356230) and maintained with daily NIM changes until D15. On D16, NIM was exchanged for retinal differentiation medium (RDM) containing DMEM (Gibco, #15140122) and DMEM/F12 (1:1), 2% B27 supplement (without vitamin A, Gibco#12587-010), NEAAs, and 1% penicillin/streptomycin (Gibco, #15140122). Feeding was done daily until D27. On D28, EBs were dislodged from the plate by checkerboard scraping, using a 10µl or 200µl pipette tip. Aggregates were washed three times and then maintained in 6 well culture plate (Greiner bio-one, #657185) in RDM medium changed every 2-3 days until D41. During this time, hRetOrg were then sorted based on their morphology under a stereoscope with transmitted light. Retinal organoids were clearly identified based on the presence of a phase-bright, pseudo-stratified neuroblastic epithelium. At D42, the medium was changed to RDM plus, consisting of RDM supplemented with 10% FBS (Gibco, #16140071), 100µM Taurine (Sigma, #T0625) and 2mM GlutaMax (Gibco, #35050061). Feeding was done every 2-3 days, depending on the color of the medium.

### Electroporation of hRetOrg

D44 through D50 hRetOrg were electroporated with a plasmid mixture of i) VSX2-EN1 (750ng/µl, final concentration) plus pUbC-mCh (150ng/µl, final concentration); ii) VSX2-EN2 (750ng/µl, final concentration) plus pUbC-mCh (150ng/µl, final concentration); iii) Stagia3 vector (750ng/µl, final concentration) plus pUbC-mCh (150ng/µl, final concentration) as previously described (Billings et al, da Silva and Cepko). Briefly, electroporations were conducted with a NEPA21 type II Nepagene electroporator using a home-designed electroporation chamber dedicated for Organoids, fitting maximum of 5 organoids at the time. 5 pulses of 40V and 50msec pulse length with 950msec interpulse interval were applied. After electroporation organoids were returned to the incubator for additional 2 days. While in culture, organoids were imaged under EVOS M5000 (Thermofisher) equipped with fluorescence filters for GFP and TX Red to check for electroporation efficiency and putative enhancer-driven GFP signal.

### Histochemical AP staining of whole hRetOrg

2 days after electroporation, hRetOrg were fixed with 4% PFA at RT for 15min, washed 3 times with 1X PBS and processed as whole for AP staining. Briefly, endogenous alkaline phosphates were heat-inactivated by incubation at 68°C for 1h15min in a water bath. Organoids were cooled down to RT for 15min, pre-incubated in NTM, pH9.5 (0.1M Tris-HCl, pH9.5, 0.1M NaCl, 0.05M MgCl_2_) for 10min and AP signal was revealed by incubation in NTM with NBT/BCIP substrate (Sigma, cat# 11681451001) until desired color was developed. All conditions were developed in parallel and for the same amount of time for fair comparisons.

### Immunohistochemistry

#### Mouse developing retina

The immunostaining on the mouse developing eye (E14.5) was performed as previously described [5, 92]. Briefly, wild-type and mutant eyes were dissected and fixed in 4% formaldehyde overnight in 4°C, followed by a PBS wash for 5 minutes. The tissues were subsequently subjected to graded concentrations of sucrose (5%, 10%, 20% and 40%) in 1X PBS (pH=7.4) and then cryoembedded in a mixture of 40% sucrose and OCT (Optimum Cutting Temperature) in 1:1 (v/v) ratio, and later cryosectioned at 14-16 µm thickness. Retinae were washed three times with PBST (0.5% Triton-X), blocked with 1% BSA in 1X PBS for 1hr and then incubated in primary antibody overnight at 4℃. The sections were washed 3 × 5 mins with 1X PBS and then incubated in fluorescent dye-conjugated secondary antibody for 1 hour at room temperature. After labelling, sections were incubated in 2 µg/ml Hoechst 33342 stain (Invitrogen H3570) for 10 mins, washed with 1XPBS, then mounted with ProLong Glass Antifade Mountant (Invitrogen P36980). Images were acquired on Olympus Fluoview Confocal microscope (v 4.2). Primary antibodies are listed in Table S1.

#### hRetOrg

2 days after electroporation, hRetOrg were fixed with 4% PFA at RT for 30min, washed 3 times with 1X PBS, cryo-protected with 30% sucrose, OCT-embedded, serially sectioned at 16µm, and stored at -80oC until further analysis. On the day of immunostaining, slides were warmed to RT for a few minutes, washed 3 times with PBS, blocked for 1 hour at RT in 5% Donkey Serum and 0.1% Triton X-100 in PBS. Organoids were then incubated with primary antibodies diluted in PBS O/N at 4°C. The primary antibodies used included anti-Rx (Table S1). Organoids were then washed 4 times with 1x PBS and once with 0.1% Triton X-100 in PBS, 5min each, and incubated with appropriate secondary antibody (Donkey anti-species A647, 1:500 dilution, ThermoFisher) and counterstained with Hoechst (ThermoFisher, cat#62249; 1:1000 dilution) in 1x PBS for 1h at RT, protected from light. This was followed by 3 washes with 1x PBS, 5min each. Slides were mounted with Fluoromount-G (Invitrogen, cat# 00495952) and imaged with Olympus FV12000 confocal microscope.

## Acknowledgments

We would like to acknowledge the Health Sciences Genomics Research Cores at the University of Pittsburgh. We would like to thank Jeff Gross for reading this manuscript. This work was funded by the Research to Prevent Blindness (RPB) career development award (to I.A), NIH R01 (EY030861) and (EY035619) (to I.A), R01HG003988 (to L.A.P.), Hillman Fund (to I. A), and a University of Pittsburgh start-up fund (to I. A). This research was also supported in part by the University of Pittsburgh Center for Research Computing through the resources provided, NIH Core Grant P30 (EY08098) to the Department of Ophthalmology and the Eye and Ear Foundation of Pittsburgh, and from an unrestricted grant from RBP, New York, NY. The research of L.A.P. was conducted at the E.O. Lawrence Berkeley National Laboratory under the US Department of Energy Contract DE-AC02-05CH11231, University of California.

## Data and Code availability

The sequencing data have been deposited to NCBI Gene Expression Omnibus (GEO) database with accession GSE281033, with GSE281030 for ATAC-seq data, GSE281031 for CUT & RUN seq data, and GSE281032 for RNA-seq data.

Data processing and analysis scripts can be found at https://github.com/AHK42/Sox2-VSX2_Analysis

